# Developmental Phase Transitions in Spatial Organization of Spontaneous Activity in Postnatal Barrel Cortex Layer 4

**DOI:** 10.1101/2020.05.26.117713

**Authors:** Shingo Nakazawa, Yumiko Yoshimura, Masahiro Takagi, Hidenobu Mizuno, Takuji Iwasato

## Abstract

Spatially-organized spontaneous activity is a characteristic feature of developing mammalian sensory systems. However, the transitions of spontaneous-activity spatial organization during development and related mechanisms remain largely unknown. We reported previously that layer 4 (L4) glutamatergic neurons in the barrel cortex exhibit spontaneous activity with a patchwork-type pattern at postnatal day 5 (P5), which is during barrel formation. In the current work, we revealed that spontaneous activity in barrel-cortex L4 glutamatergic neurons exhibits at least three phases during the first two weeks of postnatal development. Phase I activity has a patchwork-type pattern and is observed not only at P5, but also P1, prior to barrel formation. Phase II is found at P9, by which time barrel formation is completed, and exhibits broadly synchronized activity across barrel borders. Phase III emerges around P11 when L4-neuron activity is desynchronized. The Phase I activity, but not Phase II or III activity, is blocked by thalamic inhibition, demonstrating that the Phase I to II transition is associated with loss of thalamic dependency. Dominant-negative Rac1 expression in L4 neurons hampers the Phase II to III transition. It also suppresses developmental increases in spine density and excitatory synapses of L4 neurons in the second postnatal week, suggesting that Rac1-mediated synapse maturation could underlie the Phase II to III transition. Our findings revealed the presence of distinct mechanisms for Phase I to II and Phase II to III transition. They also highlighted the role of a small GTPase in the developmental desynchronization of cortical spontaneous activity.

**Significant statement:** Developing neocortex exhibits spatially-organized spontaneous activity, which plays a critical role in cortical circuit development. The features of spontaneous-activity spatial organization and the mechanisms underlying its changes during development remain largely unknown. In the present study, using two-photon in vivo imaging, we revealed three phases (Phase I, II, and III) of spontaneous activity in barrel-cortex layer 4 (L4) glutamatergic neurons during the first two postnatal weeks. We also demonstrated the presence of distinct mechanisms underlying phase transitions. Phase I to II shift arose from the switch in the L4-neuron driving source, and Phase II to III transition relied on L4-neuron Rac1 activity. These results provide new insights into the principles of developmental transitions of neocortical spontaneous-activity spatial patterns.

## Introduction

Synchronized spontaneous activity is a hallmark of mammalian sensory systems during early postnatal stages. It may support Hebbian-type synaptic competition to instruct neuronal circuit self-organization^1-5^. The spontaneous activity of the developing brain is extensively studied in the mammalian visual system. Wave-type propagation of spontaneous activity is observed in cultured retinas prepared from animals before eye opening^6^. The retinal wave is transmitted to the visual cortex through the thalamus as well as to the superior colliculus *in vivo*^7-9^. Accumulating evidence suggests a critical role for this patterned activity in visual circuit refinement^10-12^. Spontaneous activity with unique spatial organization has also recently been identified and characterized in other sensory systems of neonatal rodents^13-18^. We previously found a patchwork-type pattern in spontaneous activity of the neonatal mouse barrel cortex^16^. In that study, we generated thalamocortical axon (TCA)-RFP transgenic mice exhibiting labeling of the barrel map, to precisely identify the barrel-cortex L4 *in vivo*. L4 glutamatergic neurons were labeled in these mice with GCaMP6s via *in utero* electroporation (IUE). Taking advantage of these two methods, we were able to analyze the spatial pattern of L4-neuron spontaneous activity in relation to the barrel map. L4 neurons within the same barrel fire together in the absence of sensory input at postnatal day 5 (P5), generating a barrel-corresponding patchwork-type pattern^16^. At P11–P13, L4 glutamatergic neurons showed sparse spontaneous firing with no persisting patchwork-type patterns^16^.

The current study extended our previous work and characterized the nature of spontaneous activity in L4 glutamatergic neurons of the barrel cortex in detail. We first asked the following two questions: (1) how is spontaneous activity in barrel-cortex L4 spatially organized during the developmental stage prior to barrel map formation? If the patchwork-type pattern of activity is important for barrel map formation, the observation of similar activity is expected prior to map formation; and (2) how does spontaneous-activity transition from patchwork-type activity at P5 to sparse-type activity at P11–P13? To address these questions, we conducted *in vivo* two-photon calcium imaging of L4 glutamatergic neurons in the mouse barrel cortex at several time points during the first two weeks of postnatal development. We found that at P1, even before the initiation of barrel formation, L4-neuron spontaneous activity showed a patchwork-type pattern. The presence of a novel type of spontaneous L4-neuron activity with a spatial pattern distinct from the patchwork-type and sparse-type ones was observed at P9. These observations suggest that L4-neuron spontaneous activity exhibits at least three phases during the first two weeks of postnatal development. Phase I included P1–P5 and exhibited a patchwork-type pattern. Phase II was detected around P9 and showed wide-area synchronization. Phase III included P11–P13 and showed sparse firing. Second, we investigated the mechanism underlying the Phase I to II transition. We found that Phase I activity, but not that of Phase II or III, depended on thalamocortical input. Thus, the Phase I to II transition was characterized by a shift in the activity source. Finally, we investigated the mechanism underlying the Phase II to III transition. We revealed that this transition required L4-neuron Rac1 activity. We also found that Rac1 activity was involved in the developmental increase of excitatory synapses on L4 neurons, which may underlie the Phase II to III transition. Thus, the results of this study provide dynamic and mechanistic insights into the spatial organization of spontaneous network activity during postnatal neuronal circuit maturation.

## Results

### Patchwork-Type Pattern of L4-Neuron Spontaneous Activity Prior to Barrel Formation

We previously reported that spontaneous activity in barrel-cortex L4 shows barrel-corresponding patchwork pattern at P5 during the barrel formation period^16^. If this pattern of spontaneous activity plays an important role in barrel circuit maturation, it is expected that similar activity would be observed prior to barrel formation. To examine this possibility, GCaMP6s was transfected into L4 glutamatergic neurons of the barrel cortex by IUE and calcium transients were acquired *in vivo* by two-photon microscopy at P1 and P3 (**Fig. 1a**). P5 mice were also analyzed as a known control. We found that, at P1 and P3, L4 neurons in the barrel cortex showed spontaneous activity with a similar spatial pattern to L4 neurons at P5 (**Fig. 1b; Mov. 1**). When regions of interest (ROIs) were uniformly placed in the barrel field, a group of ROIs located nearby showed high pairwise correlation with each other but low correlation with ROIs in different locations (**Fig. 1c, d**). Although the barrels are not visible at P1, it is likely that the spatial organization of the spontaneous activity found at P1 corresponds to the prospective barrel map. To characterize the spatial organization of L4-neuron spontaneous activity at P1 in more detail, we defined an active contour as the region where GCaMP fluorescence was over the threshold in a timeframe (see Methods for details). An active contour at P5 largely corresponded to a barrel, although sometimes it corresponded to two or more barrels when neighboring barrels fired together. We found that the active contour size at P1 was slightly larger than that at P5 (**Fig. 1e**). Consistent with this result, at P1, neighboring contours showed some level of overlap (**Fig. 1f**). ROIs located on these overlapping regions showed high correlation coefficients with ROIs on each contour (**Fig. 1d**). These features of the activity pattern at P1 may reflect the fact that at this age TCA termini corresponding to different whiskers are not yet segregated. The patchwork pattern could be refined at the same time as barrel formation during neonatal development. Taken together, these results suggest that L4-neuron spontaneous activity shows a patchwork-type pattern immediately after birth even before barrel formation, although the early patchwork pattern exhibits some distinct characteristics from the one at P5.

L4-neuron spontaneous activity at P5 is blocked by whisker-pad local anesthesia, suggesting that the activity originates from the periphery^16^. To test whether P1 activity also originates from the periphery, we used the local anesthetic lidocaine in P1 mice (**Fig. 1g**). We found that lidocaine injection into the whisker pad abolished L4-neuron activity in the contralateral barrel cortex at P1 (**Fig. 1h**). These results suggest that the periphery is the source of spontaneous activity in barrel-cortex L4 at P1, similar to P5.

**Figure 1.**
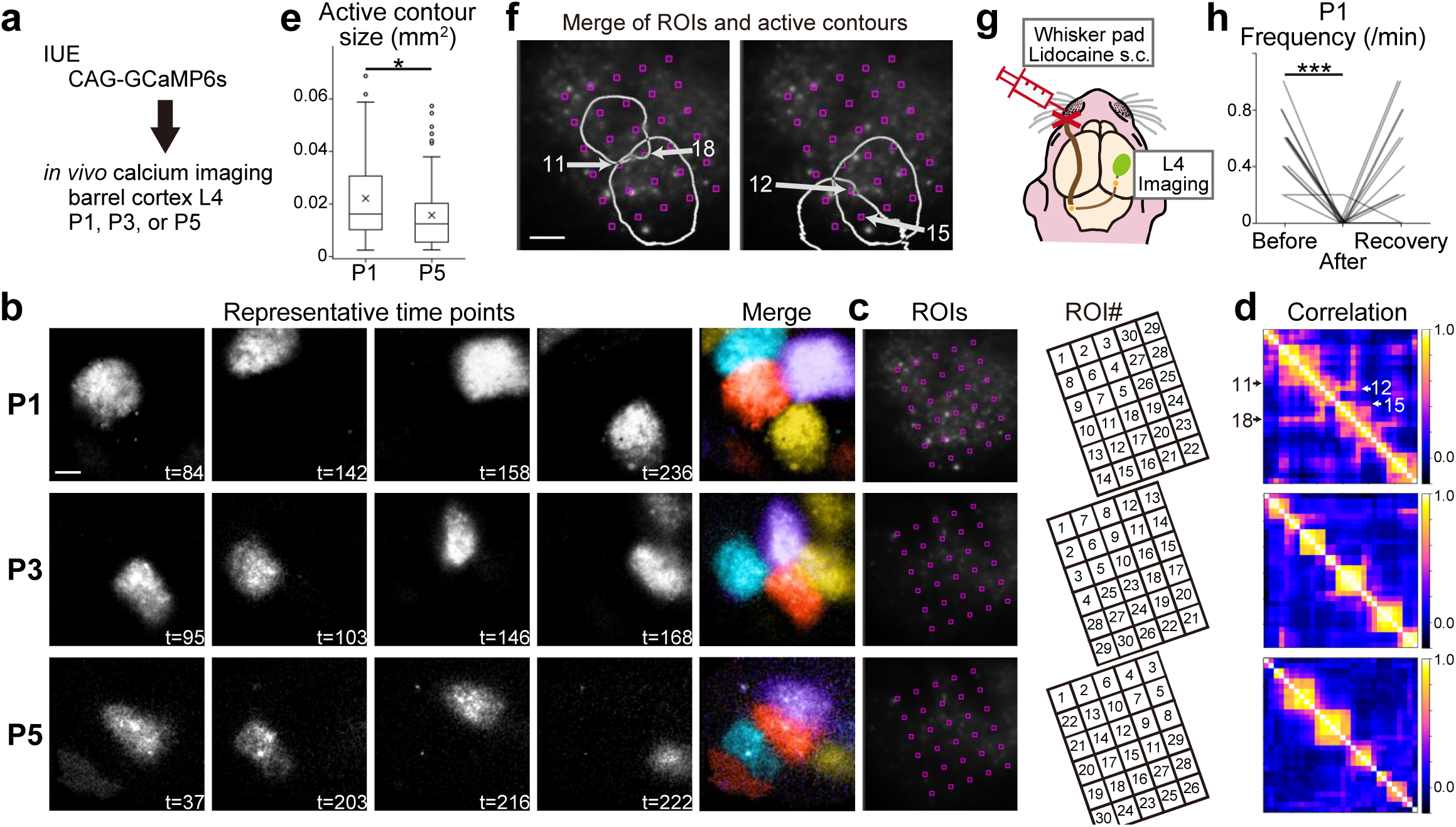
Patchwork-type spontaneous activity observed in L4 glutamatergic neurons of the barrel cortex in the first week of postnatal development. **a**, Schematic of *in vivo* imaging with dense L4-neuron labeling. CAG-GCaMP6s plasmid vector was transfected by *in utero* electroporation (IUE). **b**, Representative examples of GCaMP6s signals at P1, P3, and P5. Brightness was adjusted for visualization. See also **Mov. 1**. t: time (sec). **c**, (Left) ROIs were placed in the barrel field uniformly, because barrels are not visible at P1 yet. (Right) ROIs were manually numbered to show the correlation clusters of ROIs in **d** clearly. **d**, Correlation matrices of ROI pairs in **c. e**, Size of active contours at P1 was larger than that at P5. p = 0.011, t = 2.564, g = 0.418. n = 76 contours/3 mice at P1, 73 contours/3 mice at P5. Box plot interpretation is described in the Methods. **f**, Two examples of overlapping active contours at P1. Arrows indicate ROIs located on overlapping regions of neighboring contours. ROI numbers are the same as those shown in **c. g**, Schematic of the L4-neuron imaging experiment with peripheral silencing. **h**, Local anesthetic lidocaine injection into the whisker pad significantly reduced the frequency of L4-neuron spontaneous activity in contralateral side of the barrel cortex at P1 (Before vs After: p < 0.001, t = 8.101, g = 3.177; n = 13 ROIs/2 mice). Note that the activity was recovered in 15 minutes (Recovery), suggesting that lidocaine injection did not damage the circuits. (**b, f**) Scale bars, 100 μm.

### Wide-Area Synchronization of L4-Neuron Spontaneous Activity at P9

To investigate how spontaneous network activity transitions from patchwork-type at P1–P5 to sparse-type at P11–P13, we analyzed the spatial organization of L4-neuron spontaneous activity at P9. We first compared the activity pattern between P9 and P5 by transfecting GCaMP6s into a dense population of L4 glutamatergic neurons and performing calcium imaging *in vivo* (**Fig. 2a**). ROIs were placed on barrels that were visualized using TCA-RFP Tg mice or *post-hoc* staining with an anti-VGluT2 antibody and/or DAPI (**Fig. 2b**). At P5, L4 neurons showed spontaneous activity corresponding to the barrel map (**Fig. 2c, d**) as previously reported^16^. ROIs put on the same barrels tended to fire together (**Fig. 2c**). Therefore, ROI pairs in the same barrels were highly correlated with each other, while ROIs located in different barrels tended to fire independently and showed low correlations (**Fig. 2d–f**). While, at P9, L4-neuron spontaneous activity was widely synchronized in the barrel cortex across the barrel borders (**Fig. 2c, d; Mov. 2**). They were still highly correlated even at large distance between ROI pairs (**Fig. 2e**) and regardless of whether ROI pairs were in the same barrel or not (**Fig. 2f**). Overall, L4 neurons at P9 showed much higher correlations than those at P5 (**Fig. 2g**). Thus, the spatial pattern of L4-neuron spontaneous activity at P9 was distinct from that at P5.

**Figure 2.**
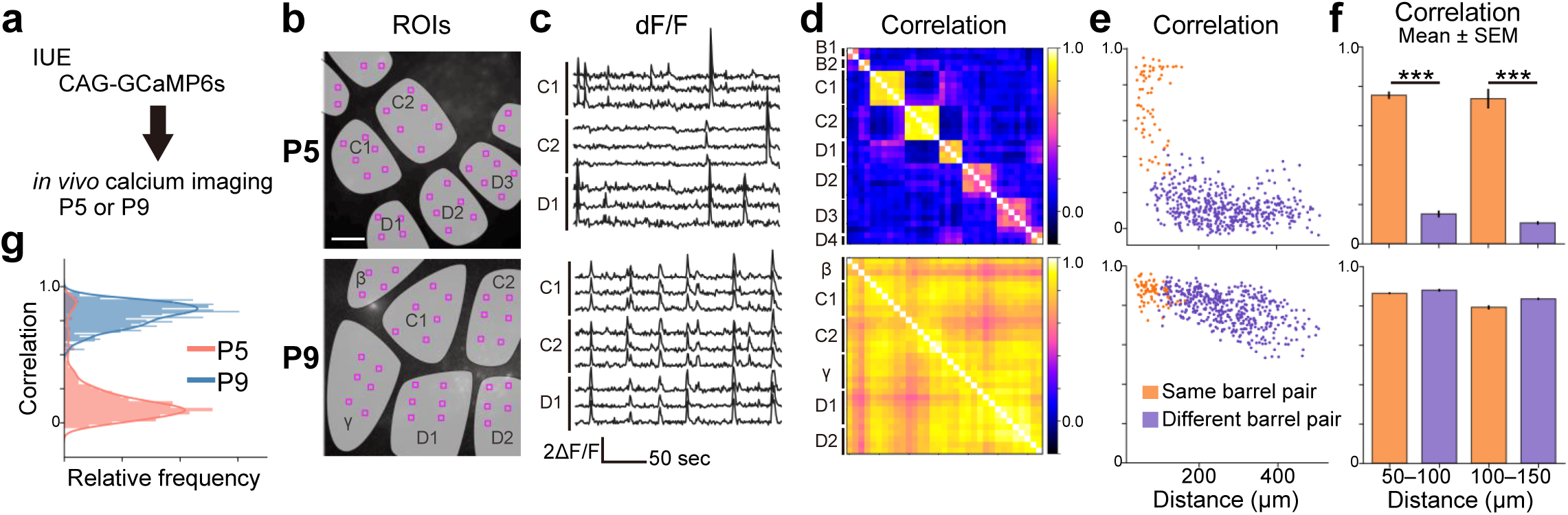
Spatial organization of L4-neuron spontaneous activity at P9 is different from that at P5. **a**, Schematic of *in vivo* imaging with dense L4-neuron labeling. **b**, ROIs were placed on each barrel. Scale bar, 100 μm. **c**, Representative examples of *in vivo* calcium transients at P5 and P9. See also **Mov. 2. d**, Pairwise ROI correlation matrices. **e**, Pairwise ROI correlation as a function of pair distance. **f**, Comparison of correlation coefficients between ROI pairs from the same barrel (orange) and from different barrels (purple). (Top) At P5, ROI pairs from the same barrels were highly correlated, and those located in different barrels showed low correlations even the distance between ROI pairs were similar (50–100 μm: p < 0.001, t = 23.024, g = 3.798, n = 84 same barrel pairs and 39 different pairs; 100–150 μm: p < 0.001, t = 12.369, g = 4.842, n = 21 same and 128 different; 2 mice). (Bottom) At P9, ROI pairs were highly correlated regardless of whether they were in the same barrel or not (50–100 μm: n = 151 same and 58 different pairs; 100–150 μm: 50 same and 175 different paris; 3 mice). **g**, Distribution of pairwise correlation coefficients at P5 and P9 shown in **d**. P9 activity showed much higher synchrony than P5 activity.

We next compared the P9 and P11–P12 activity patterns by using the Supernova method^19,20^ to label a small population of L4 neurons with GCaMP6s and conducting *in vivo* calcium imaging in single-cell resolution (**Fig. 3a**). We previously reported that L4 neurons show asynchronous patterns of spontaneous activity at P11–P13^16^. Here, we confirmed that L4 neurons fire sparsely and show no clear spatial organization of spontaneous activity at P11–P12. At this age, pairwise correlation coefficients were low between ROIs placed on individual L4 neurons (**Fig. 3b–d**) and synchronized firing involving many neurons was not observed (**Fig. 3e**). In contrast, at P9, as observed in dense cell-labeling experiments (**Fig. 2)**, L4 neurons showed high correlations even when they were in different barrels (**Fig. 3b–d**) and tended to fire synchronously (**Fig. 3e**). Overall, L4 neurons at P9 showed much higher correlations than those at P12 (**Fig. 3f**). Quantitative analyses revealed that the frequency of neuronal firing events was similar between P9 and P11– P12 (**Fig. 3g**). However, the ratio of synchronously firing events to total firing events was significantly higher at P9 than at P11–P12 (**Fig. 3h**). Further, the ratio of neurons that participated in individual synchronized events to total neurons with ROIs was significantly higher at P9 (72 ± 5%, Mean ± SD) than P11–P12 (26 ± 3%) (**Fig. 3i**). These results indicate that a large population of neurons distributed over multiple barrels fired together at P9. Thus, P9-type L4-neuron spontaneous activity is distinct from P11–P13-type activity.

**Figure 3.**
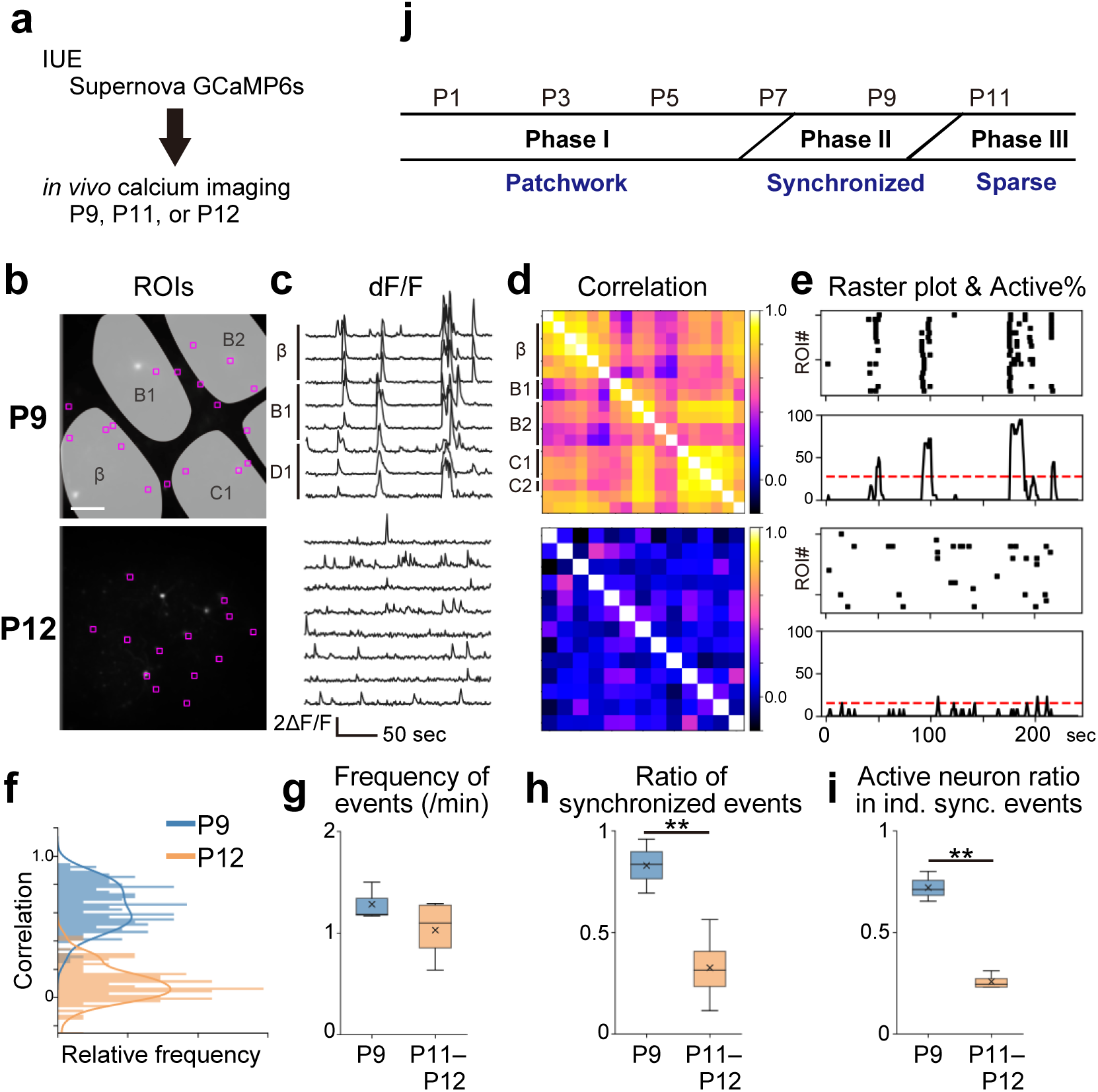
Spatial organizations of L4-neuron spontaneous activity in the barrel cortex at P9 and P11–P12. **a**, Schematic of *in vivo* calcium imaging with Supernova-mediated sparse L4-neuron GCaMP6s labeling. **b**, ROIs were placed on individual L4 neurons. Scale bar, 100 μm. **c**, Representative examples of *in vivo* calcium transients. **d**, Pairwise ROI correlation matrices. **e**, Raster plots and activity histograms. Dashed red lines indicate the chance rate (p = 0.01). **f**, Distribution of pairwise correlation coefficients shown in **d. g**, Frequency of neuronal firing events. ROI averages of individual mice at P9 and P11–P12 were compared (p = 0.239, t = 1.340, g = 0.945). **h**, The ratio of synchronized events to total firing events. ROI averages for individual mice were compared between P9 and P11–P12 (p = 0.009, t = 4.170, g = 3.012). **i**, The ratio of active ROIs to total ROIs in individual synchronized events. Maximum number of ROIs that fired together during individual synchronized events was divided by total ROI number, and averages for all synchronized events in individual mice were compared between P9 and P11-P12 (p = 0.003, t = 9.901, g = 8.378). n = 3 (P9) and 4 (P11–P12) mice. **j**, Three phases of spontaneous network activity observed in barrel-cortex L4 neurons during the first two weeks of postnatal development.

These results suggest that there are at least three phases of spatial organization in L4-glutamatergic-neuron spontaneous activity during the first two weeks of postnatal development (**Fig. 3j**). Phase I is around P1 to P5 and shows a patchwork-type pattern. Phase II is around P9 and shows widely synchronized activity across the barrel borders. Phase III is around P11–P12 and shows sparse firing.

### Phase II and Phase III Activity is Independent of Thalamocortical Inputs

We previously reported that Phase I spontaneous activity is transmitted to L4 via TCAs^16^. To observe whether Phase II and Phase III activity depends on TCA input, we expressed the inhibitory DREADD hM4Di in the sensory thalamus by mating 5HTT-Cre Tg^21^ and Cre-dependent hM4Di^22^ mice. IUE was used to transfect L4 neurons of these mice with GCaMP6s. *In vivo* calcium transients in barrel-cortex L4 neurons were analyzed before and after clozapine-N-oxide (CNO) administration (**Fig. 4a**).

**Figure 4.**
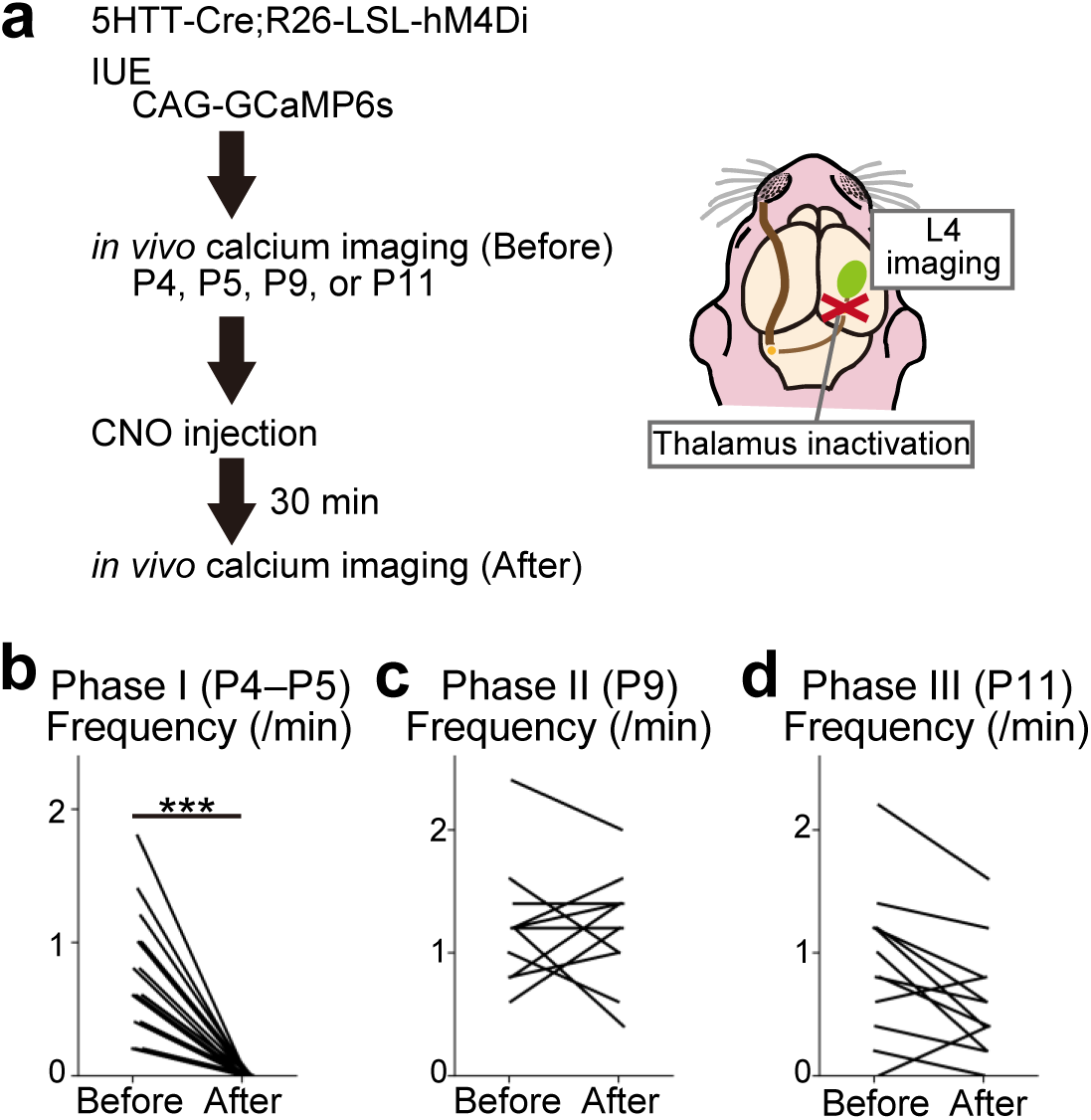
Thalamic inactivation silences L4-neuron spontaneous activity in Phase I but not in Phase II or Phase III. **a**, Schematics of *in vivo* calcium imaging of L4 neurons before and after the inhibitory DREADD-mediated thalamus silencing. CAG-GCaMP6s plasmid was transfected to 5HTT-Cre;R26-loxP-STOP-loxP (LSL)-hM4Di mice by IUE. **b**, Silencing thalamic neurons significantly reduced the frequency of L4-neuron spontaneous activity in Phase I. p < 0.001, t = 7.400, g = 2.283. n = 21 ROIs/3 mice. **c–d**, Silencing the thalamic neurons had no significant effect on frequency of L4-neuron spontaneous activity in both Phase II (**c**: p = 0.928, t = 0.091, g = 0.039. n = 11 ROIs/2 mice) and Phase III (**d**: p = 0.156, t = 1.471, g = 0.601. n = 12 ROIs/2 mice).

Before analyzing Phase II and Phase III activity, we examined whether the inhibitory DREADD system is effective in the developing thalamus, by analyzing Phase I (P4–P5) as a control. We conducted *in vivo* two-photon calcium imaging of the barrel-cortex L4 before and after CNO administration at P4 or P5. We found that L4 spontaneous activity was silenced following CNO administration (**Fig. 4b**), confirming that Phase I activity is dependent upon the thalamus. This result also suggests that thalamic expression of inhibitory DREADDs effectively blocked thalamic activity in early postnatal stages. Therefore, we conduced calcium imaging of thalamic-inhibitory-DREADD mice at P9 (Phase II) and P11 (Phase III). L4-neuron spontaneous activity was not significantly changed between before and after CNO application (**Fig. 4c, d**). These results suggest that Phase II and Phase III activity is independent of thalamic inputs.

In conclusion, Phase I activity arises from the periphery and is transmitted to the cortex via TCAs. Conversely, Phase II and Phase III activity occurs independent of thalamic inputs. Thus, the transition from Phase I to Phase II is associated with a switch in the source of spontaneous activity.

### Dominant-Negative Rac1 Suppresses Increases in Spine Density During the Second Week of Postnatal Development

We next investigated the mechanism that controls the transition of Phase II to Phase III L4-glutamatergic-neuron spontaneous activity in the barrel cortex. Although sparse spontaneous activity is observed in various areas, layers and cell types of the mammalian cortex in the late developmental stage^14,15,23-25^, the underlying mechanisms of sparsification remain largely unresolved. Notably, little attention has been paid to the possible involvement of maturation of the excitatory system. Given that dendritic spine density dramatically increases in the second week of postnatal development^26-29^, maturation of excitatory circuits may affect sparsification of L4-glutamatergic-neuron spontaneous activity. If this hypothesis is correct, blocking developmental spinogenesis could inhibit sparsification of L4-neuron spontaneous activity.

To test this hypothesis, we first examined whether spine density increased in barrel-cortex L4 glutamatergic neurons during the transition from Phase II to III. To reduce data dispersion, we focused our analyses on barrel-inner dendrites of barrel-edge spiny stellate neurons, which are the major type of L4 glutamatergic neurons in the barrel cortex (see Methods). We found that spine density more than doubled between P9 and P11, with increases in the mushroom- and thin/stubby-types (**Fig. 5a–c**).

**Figure 5.**
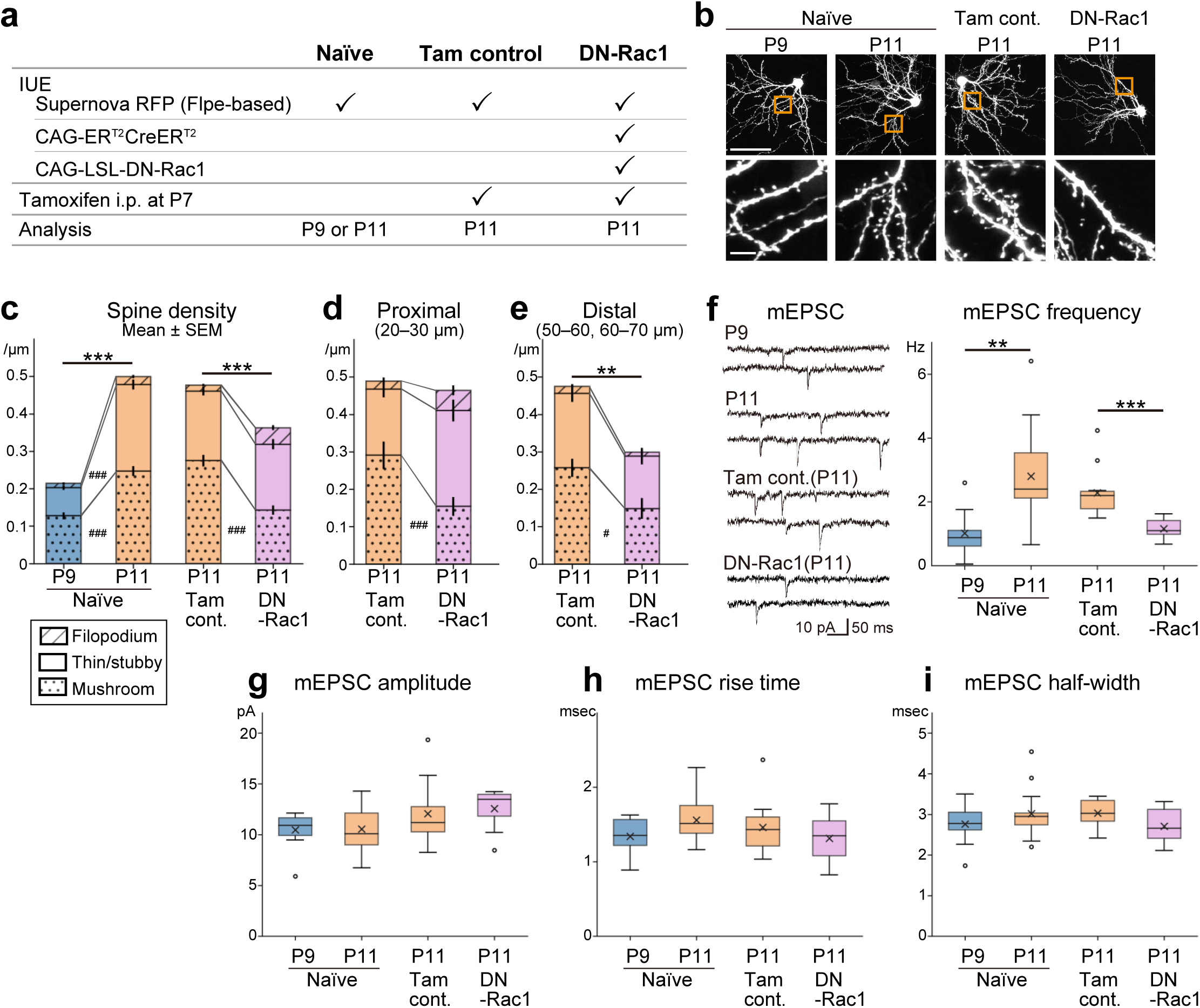
Dendritic spine maturation between Phase II and Phase III periods were interfered by dominant-negative (DN)-Rac1 expression in L4 neurons. **a**, Schematic of spine analysis experiments. L4 glutamatergic neurons were sparsely labeled with flpe-based Supernova RFP. For DN-Rac1, CAG-ER^T2^CreER^T2^ and CAG-LSL-DN-Rac1 plasmids were also transfected. For Tam control and DN-Rac1 mice, tamoxifen was administrated by intraperitoneal (i.p.) injection at P7. **b**, Representative examples of L4 spiny stellate neurons of naïve P9, naïve P11, Tam control P11, and DN-Rac1 P11. Scale bar, 50 μm (top) and 5 μm (bottom). **c**, Spine density of barrel-cortex L4 spiny stellate neurons of naïve P11 was significantly higher than that of naïve P9 (p < 0.001, t = 11.770, g = 1.533). Mushroom-type (p < 0.001, t = 7.361, g = 0.970) and thin/stubby-type (p < 0.001, t = 10.488, g = 1.362) spines increased from P9 to P11 (Welch’s t-test with Holm correction. n = 102 (P9) and 117 (P11) segments). DN-Rac1 neurons had significantly less spine density than Tam control neurons at P11 (p < 0.001, t = 4.308, g = 0.564). Mushroom-type spine density was also lower in DN than in Tam (p < 0.001, t = 6.647, g = 0.840) (Welch’s t-test with Holm correction. n = 134 (Tam) and 94 (DN) segments). **d**, On the proximal dendrites (20–30 μm from soma center), total spine density of DN neurons was similar to that of Tam control at the same age (P11). However, mushroom-type spine density was lower in DN (p = 0.004, t = 3.068, g = 0.911. n = 23 (Tam) and 20 (DN) segments). **e**, On the distal dendrites (50–60, 60–70 μm), total spine density of DN was lower than that of Tam control (p = 0.002, t = 3.453, g = 0.829), and mushroom-type spine density was also significantly lower in DN (p = 0.005, t = 2.922, g =0.729) (Welch’s t-test with Holm correction, n = 42 (Tam) and 21 (DN) segments). **c–e**, Data were obtained from 3, 3, 4, 3 neurons of 3, 3, 3, 3 naïve P9, naïve P11, Tam P11, DN P11 mice, respectively. **f**, The average frequency of miniature EPSCs (mEPSCs) was significantly increased from P9 to P11 in naïve animals (p = 0.002, t = 3.633, g = 1.396). By overexpressing DN-Rac1, the average frequency of mEPSCs became lower than that of Tam control (p < 0.001, t = 4.522, g = 1.809). The same experimental scheme shown in (**a**) was used for Tam control and DN-Rac1 of the physiological analysis. **g–i**, Additional properties of mEPSCs. The amplitude (**g**, p = 0.899, t = 0.129, g = 0.053), rise time (**h**, p = 0.065, t =1.951, g = 0.802), and half-width (**i**, p = 0.298, t = 1.067, g = 0.434) of mEPSCs did not significantly change from P9 and P11 in naïve mice, and also no significant differences between Tam control and DN-Rac1 (p = 0.654, t = 0.455, g = 0.188 for amplitude; p = 0.327, t = 1.005, g = 0.426 for rise time; p = 0.075, t = 1.900, g = 0.835 for half-width). **f–i**, Data were obtained from 10, 13, 12, 10 neurons of 4, 4, 4, 5 naïve P9, naïve P11, Tam P11, DN P11 mice, respectively.

We used Rac1 (T17N), a dominant-negative (DN)-Rac1 that blocks spinogenesis in cultured hippocampal and cortical neurons^30,31^, to interfere with developmental spinogenesis. We used an inducible gene expression strategy to overexpress DN-Rac1 because constitutive DN-Rac1 expression interferes with radial migration of transfected cells^32^. IUE was used to transfect L4 glutamatergic neurons with tamoxifen-inducible Cre (ER^T2^CreER^T2^)^33-35^ and Cre-dependent DN-Rac1. Flpe-based Supernova RFP^19,20^ was co-transfected for clear visualization of neuronal morphology. Then, tamoxifen was intraperitoneally injected at P7 and brain samples were collected at P11 (**Fig. 5a**). Comparing DN-Rac1 overexpression and tamoxifen control (Tam control) at P11 indicated that DN-Rac1 overexpression decreased spine densities (**Fig. 5b, c**). In DN-Rac1-expressing neurons, mushroom-type spine density was also lower than in Tam control neurons (**Fig. 5c**). This effect was more pronounced on the distal dendrites than in the proximal dendrites. On the proximal dendrites, mushroom-type spine density was decreased, but the total spine density was not changed in DN-Rac1-overexpressing neurons (**Fig. 5d**). On the distal dendrites, total spine and mushroom-type spine densities were decreased (**Fig. 5e**). These results suggest that DN-Rac1 overexpression inhibited increases in and maturation of dendritic spines between Phase II and Phase III.

We also assessed electrophysiological synaptic properties of DN-Rac1-overexpressing neurons. Like above, DN-Rac1 overexpression was induced by tamoxifen injection at P7, and *ex vivo* whole-cell patch-clamp recordings were conducted at P11 (**Fig. 5a**). We analyzed miniature EPSCs (mEPSCs) recorded from L4 spiny stellate neurons identified by *post-hoc* staining with biocytin. In naïve animals, the frequency of mEPSCs increased from P9 to P11 (**Fig. 5f**). We found that the frequency was lower in DN-Rac1-overexpressing cells compared to Tam controls at P11, which was consistent with our morphological analysis of spine density (**Fig. 5f**). There were no significant differences in the amplitude, rise time and half-width between them, suggesting that synaptic strength and kinetics did not change (**Fig. 5g–i**). These results suggest that Rac1 inhibition reduces functional excitatory inputs to barrel-cortex L4 neurons.

Taken together, these results suggest that Rac1 is important for increases in dendritic spine density and functional maturation of excitatory synapses at L4 glutamatergic neurons, which occur between the Phase II and Phase III periods.

### DN-Rac1 Hampers the Phase II to III Transition

Finally, we conducted *in vivo* calcium imaging of DN-Rac1-overexpressing L4 neurons. L4 neurons were co-transfected with ER^T2^CreER^T2^, Cre-dependent DN-Rac1, and Supernova-GCaMP6s. Tamoxifen was administrated at P7 and *in vivo* calcium imaging was conducted at P11 in single-cell resolution (**Fig. 6a**). As expected, Tam controls showed sparse firing (**Fig. 6b–d**). Notably, we observed widely synchronized activity across the barrel borders in DN-Rac1 mice (**Fig. 6b, c; Mov. 3**). Synchronized events were frequently observed in the activity histogram of DN-Rac1 mice (**Fig. 6d**). Quantitative analyses showed that the frequency of neuronal firing events was similar between Tam control and DN-Rac1 mice (**Fig. 6e**). However, the ratio of synchronized events to total events was significantly higher in DN-Rac1 compared to Tam controls (**Fig. 6f**). These results indicate that the synchronized events were prominent in DN-Rac1 cases. The ratio of active neurons to total neurons in individual synchronized events tended to be higher in DN-Rac1 than in Tam controls although it was not significantly different (**Fig. 6g**). These features of L4-neuron spontaneous activity in DN-Rac1 mice at P11 were distinct from Phase III-type sparse firing in naïve P11 mice and appeared similar to Phase II-type spontaneous activity (**Fig. 3**). However, it should be noted that DN-Rac1 activity at P11 was not identical to Phase II activity observed in naïve P9 mice. For example, the ratio of synchronized events to total firing events was 83 ± 9% (mean ± SD) for naïve mice at P9 (**Fig. 3h**), but 63 ± 13% for DN-Rac1 mice at P11 (**Fig. 6f**). Also, the ratio of neurons involved in each synchronized event was 72 ± 5% for naïve mice at P9 (**Fig. 3i**), but 53 ± 17% for DN-Rac1 mice at P11–P12 (**Fig. 6g**). IUE-mediated transfection, tamoxifen-induced Cre recombination, or DN-Rac1 effects may be insufficient to stop the phase transition completely. These results suggest that DN-Rac1 overexpression partially blocked the sparsification process that occurs during normal development between P9 to P11.

**Figure 6.**
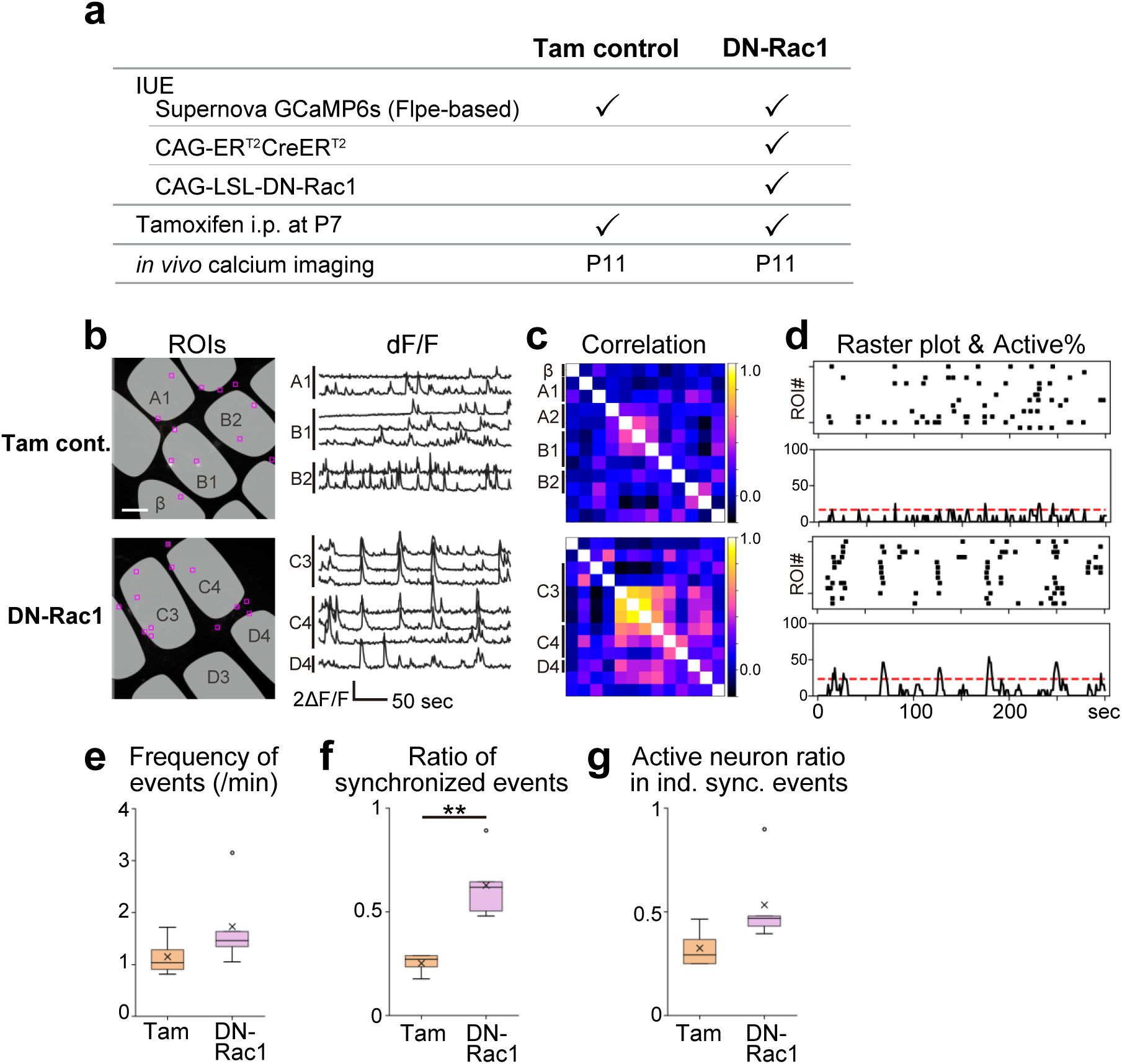
The Phase II to III transition was affected by DN-Rac1 overexpression in L4 glutamatergic neurons. **a**, Schematic of experiments. For DN-Rac1, Flpe-based Supernova-GCaMP6s, CAG-ER^T2^CreER^T2^ and CAG-LSL-DN-Rac1 vectors were co-transfected into TCA-RFP mouse by IUE. **b**, Representative examples of *in vivo* calcium transients at P11. Scale bar, 100 μm. See also **Mov. 3. c**, Pairwise ROI correlation matrices. **d**, Raster plots and activity histograms. Dashed lines in activity histograms indicate chance rate (p = 0.01). **e**, Frequency of neuronal firing events was similar between Tam control and DN-Rac1 mice (p =0.217, t = 1.378, g = 0.855). **f**, The ratio of synchronized events (> chance rate) to total firing events was higher in DN-Rac1 than Tam control (p = 0.005, t = 4.821, g = 2.921). **g**, In average 53 ± 17% (mean ± SD) of ROIs were fired in individual synchronized events in DN-Rac1 mice, while, 32 ± 8% in Tam control (p = 0.093, t = 1.992, g =1.237). n = 4 (Tam control) and 5 (DN-Rac1) mice (**e–g**).

Taken together, these results reveal that Rac1 plays an important role in the Phase II to III transition of L4-glutamatergic-neuron spontaneous activity in the barrel cortex.

## Discussion

Spatial organizations of spontaneous activity provide a template for activity-dependent development of mammalian neuronal circuits. We showed that there are at least three phases (Phases I, II, and III) in spatial organization of barrel-cortex L4-glutamatergic-neuron spontaneous activity during the first two postnatal weeks and further demonstrated mechanisms that are involved in Phase I to II and Phase II to III transitions.

### Phase I Spontaneous Activity of L4 Glutamatergic Neurons

We previously reported that barrel-cortex L4 glutamatergic neurons exhibit patchwork-type spontaneous activity at P5^16^. Herein, we demonstrated that similar patchwork-type spontaneous activity was present at P1 (**Fig. 1**), prior to barrel map formation^20,36^. In the mouse barrel-cortex L4, thalamocortical connectivity is drastically reorganized to form the barrel map during the first postnatal week^20,37-39^. The patchwork-type spontaneous activity may play an important role in thalamocortical reorganization.

Thalamocortical slices prepared from mouse embryos exhibit a wave-type spontaneous activity that propagates in the thalamus^40,41^. On the other hand, our analysis of cortical L4-neuron activity at P1 did not indicate the presence of wave-type activity. Similar to that observed at P5^16^, L4 activity at P1 was abolished by whisker-pad lidocaine administration (**Fig. 1g, h**), suggesting that Phase I activity arises from the periphery, rather than thalamus. The thalamic wave reported in the embryonic thalamus *in vitro* may not be present or sufficiently strong to induce L4-neuron calcium transients in the postnatal brain *in vivo*.

L2/3-neuron spontaneous activity was analyzed in the rat barrel cortex at P4–P7^14^. These neurons exhibited Phase I-like activity, which was highly synchronous within local clusters. However, the spatial pattern of L2/3-neuron spontaneous activity does not track with specific barrel boundaries^14^. At this age, L4-neuron axons projecting to L2/3 are sparse and not confined within a barrel column^42^. Such rudimentary axonal projections may explain the discrepancy between L4 and L2/3. It is unclear whether the patchwork pattern observed in L4 neurons at P1 corresponds to the barrel map because the barrel map does not exist at P1. A previous *in vivo* imaging study of the rat barrel cortex showed that the spontaneous firing clusters were similar to the cortical representations of whisker stimulation at P0–P1^18^. Technical improvement of the long-term *in vivo* imaging of the neonatal cortex^38^ would enable imaging starting at P1. Such studies will determine whether patchwork-type activity at P1 corresponds to the prospective barrel map. Spontaneous activity of GABAergic neurons in the neonatal mouse barrel cortex was recently reported^24^. Further studies are required to fully understand the characteristics and interactions of Phase I-type locally clustered spontaneous activity among various cortical layers and cell-types in neonatal stages and their roles in somatosensory circuit maturation.

### Phase II Spontaneous Activity and Phase I to II Transition

During the Phase II period around P9, L4 glutamatergic neurons showed spontaneous activity that was widely synchronized across multiple barrels (**Figs. 2, 3**). The current study also found that the Phase I to II shift in L4-glutamatergic-neuron spontaneous activity arose from the switch in the L4-neuron driving source. Phase I L4-neuron activity was blocked by whisker-pad lidocaine administration and DREADD-induced thalamic inactivation (**Figs. 1, 4**), suggesting that this phase of activity originates in the periphery and is relayed to L4 via TCAs. Conversely, Phase II and III activity failed to be blocked by DREADD-induced thalamic inactivation (**Fig. 4**), suggesting that these phases of L4 activity are not relayed via TCAs.

The Phase I to II transition period is associated with thalamocortical connectivity maturation, e.g., the barrel map, representing whisker-related clusters of TCA termini, is established in the first week of postnatal development^20,36^. Long-term potentiation is easily induced at thalamocortical synapses at P3–P7, but not P9–P11 ^43^. L4 neurons drastically reorganize their dendritic projection patterns during the first postnatal week, and by P9, they establish adult-type dendritic projections^20,38,44^. Subplate neuron neurites that initially accumulate in the barrel hollow gradually shift to the barrel septa between P6 and P10^45^. The barrel net, which is the whisker-related axonal pattern of L2/3 neurons in the barrel-cortex L4, is not present at P6 but is formed by P10^46^. Thus, thalamocortical and/or cortical maturation during these periods could play an important role in terminating the prevailing role of thalamocortical input and initiating the critical involvement of cortical or other non-thalamic input in driving L4-neuron spontaneous activity. Large neuronal ensembles of spontaneous activity that are similar in appearance to the Phase II activity of L4 glutamatergic neurons were found in rat barrel-cortex L2/3 neurons between P8 and P11^14^. The slight discrepancy in the phase-shift timing may be attributed to differences in animals and/or layers. In mouse, spontaneous-activity assemblies of GABAergic neurons become wider from P4–P6 to P7– P9^24^. However, it remains unknown whether the Phase II-like activity of L2/3 and GABAergic assemblies of the barrel cortex are independent of thalamic input^14,24^.

### Phase II to III Transition of L4-Neuron Spontaneous Activity

The adult neocortex exhibits spontaneous activity that is sparse and heterogeneously distributed in space and time across the neuronal population^47,48^. The sparseness of this neuronal activity allows precise information processing^48-50^. L4 glutamatergic neurons in the barrel cortex started to show sparse spontaneous activity by P11 (**Fig. 3**). Mice begin to show active whisking and explorative behavior around P14^51,52^, which drastically increase the amount and complexity of information perceived and processed by L4 neurons. The developmental sparsification of L4 glutamatergic neurons may play a role in setting up the neural circuits that underlie adult barrel-cortical function.

We provided evidence that the small GTPase Rac1, which is a key regulator of actin dynamics, plays an important role in the Phase II to III transition, a time point at which L4-neuron spontaneous activity undergoes sparsification. At P11, L4 glutamatergic neurons showed sparse-type (Phase III) spontaneous activity (**Fig. 3**). However, in P11 mice with L4 neurons that express DN-Rac1, L4 neurons exhibited spontaneous activity with a spatial pattern that was similar to the Phase II type (**Fig. 6**). Rac1 plays an important role in broad aspects of brain development, including cell migration, axon guidance and dendritic spine formation, by regulating actin dynamics^53^. During cortical development, the timing of spontaneous activity sparsification coincides with a drastic increase in dendritic-spine density. We demonstrated that the spine densities of mouse barrel-cortex L4 glutamatergic neurons more than doubled between Phases II (P9) and III (P11) (**Fig. 5c**). L4-neuron developmental spinogenesis was suppressed by DN-Rac1 expression (**Fig. 5c**). Furthermore, mEPSC frequency was lower in DN-Rac1-expressing L4 neurons than in controls at P11 (**Fig. 5f**). These results suggest that Rac1 regulates the number of excitatory synapses on L4 glutamatergic neurons during the transition from Phase II to III. Thus, Rac1 may facilitate spontaneous-activity sparsification by promoting excitatory-network maturation in cortical L4.

Developmental sparsification of spontaneous activity is observed in the postnatal mammalian cortex across various areas, layers and cell types^14,15,23-25^. However, the mechanisms underlying sparsification remain largely unknown. For example, the whisker plucking starting at P2 does not affect the L2/3-neuron spontaneous-activity sparsification in the rat barrel cortex, suggesting that sensory input is dispensable for this process^14^. To our knowledge, we provided the first evidence that the functional blockade of a cortical molecule affects the developmental sparsification of cortical spontaneous activity.

Our results do not exclude the possibility that inhibitory-circuit maturation plays a role in spontaneous-activity developmental sparsification. Symmetric synapses, which are putative inhibitory synapses, are increased^27^, and intrinsic L2/3 excitability is decreased during the transition period^14^. Therefore, inhibitory-synapse maturation may also be involved in spontaneous-activity developmental sparsification. Previous experiments using dissociated neuronal cultures suggest that Rac1 affects GABAA receptor function by regulating its clustering and recycling^54,55^. Therefore, Rac1 may regulate L4-neuron spontaneous-activity sparsification by facilitating the maturation of both inhibitory and excitatory synapses.

Thus, these results suggest that Rac1 plays an important role in the Phase II to III transition of L4-glutamatergic-neuron spontaneous activity, possibly via the regulation of the synaptic maturation of L4 neurons.

## Materials and Methods

### Animals

All experiments were performed according to the guidelines for animal experimentation of the National Institute of Genetics and the National Institute of Physiological Sciences and were approved by their animal experimentation committees. The day at which the vaginal plug was detected was designated as embryonic day (E)0 and E19 was defined as P0. For electrophysiological analysis, timed-pregnant ICR mice were obtained from SLC Japan (for naïve) or CLEA Japan (for Tam control and DN-Rac1). Sex of newborn mice was not identified.

Used transgenic lines are as follows: TCA-RFP^16^, TCA-GFP^20^, 5HTT-Cre^21^, R26-LSL-hM4Di-DREADD (JAX stock #026219)^22^. TCA-RFP mice were backcrossed from B6C3F2 to ICR three to seven times. TCA-GFP mice were backcrossed from B6 to ICR 11 to 13 times. 5HTT-Cre and R26-LSL-hM4Di-DREADD mice were backcrossed from B6 to ICR one or two times.

### *In utero* electroporation (IUE)

IUE was conducted at E13 night or E14 morning. Timed-pregnant mice were anesthetized with an intraperitoneal injection of a combination anesthetic (medetomidine (0.3 mg/kg), midazolam (4 mg/kg), and butorphanol (5 mg/kg) in saline), or with an intraperitoneal injection of pentobarbital (50 mg/kg) in saline and isoflurane inhalation (1−3.5%). DNA solution (diluted in Milli-Q water and < 5% trypan blue (Sigma: T8154)) was injected into lateral ventricles of embryos via a pulled glass capillary (Drummond: 2-000-050), and two to five times square electric pulses (40 V; 50 ms) were delivered by tweezer-electrodes (NepaGene: CUY650P5) and an electroporator (NepaGene: CUY21SC). When a combination anesthetic was used, atipamezole (0.3 mg/kg) in saline was administered as an intraperitoneal injection after IUE.

#### Dense GCaMP labeling

pK152 (1,000 ng/μL) was used. To identify *in vivo* imaged area in sections, pK036 (5−15 ng/μL) + pK037 (1,000 ng/μL), or pK036 (5−15 ng/μL) + pK281 (1,000 ng/μL) were also transfected as a marker of IUE.

#### Sparse GCaMP labeling (Supernova GCaMP)

pK031 (5−15 ng/μL) and pK175 (1,000 ng/μL) were used. pK098 (500 ng/μL) was also transfected as a marker.

#### DN-Rac1 histology and electrophysiology

pK036 (5−15 ng/μL), pK037 (1,000 ng/μL), pK326 (200 ng/μL), and pK328 (1,000 ng/μL) were used. For Tam control, pK036 (5−15 ng/μL) and pK037 (1,000 ng/μL) were transfected.

#### DN-Rac1 calcium imaging

For DN-Rac1, Flpe-based Supernova GCaMP6s [pK036 (5−15 ng/μL), pK313 (1,000 ng/μL), pK326 (200 ng/μL), and pK328 (1,000 ng/μL)] were transfected. For Tam control, pK036 (5−15 ng/μL) and pK313 (1,000 ng/μL) were transfected. As a marker of IUE, pK302 (200 ng/μL) or pK037 (500 ng/μL) was also transfected.

### Plasmids

pK031: TRE-Cre^20^

pK036: TRE-Flpe-WPRE^19^

pK037: CAG-FRT-STOP-FRT-RFP-ires-tTA-WPRE^19^

pK098: CAG-loxP-STOP-loxP-nls-tagRFP-ires-tTA-WPRE^19^

pK152: CAG-GCaMP6s^16,56,57^

pK175: CAG-loxP-STOP-loxP-GCaMP6s-ires-tTA-WPRE^16^

pK281: CAG-FRT-STOP-FRT-CyRFP-ires-tTA-WPRE

pK302: CAG-tagBFP.

pK313: CAG-FRT-STOP-FRT-GCaMP6s-ires-tTA-WPRE.

pK326: CAG-ER^T2^CreER^T2^-WPRE.

pK328: CAG-loxP-STOP-loxP-Rac1(T17N).

pCAG-CyRFP was a gift from Ryohei Yasuda (Addgene plasmid # 84356; http://n2t.net/addgene:84356; RRID:Addgene_84356)^58^. pTagBFP-actin vector was from Evrogen (#FP174) ^59^. pCre-ER^T2^ was a gift from Piewe Chambon^34,60,61^. pCyPet-Rac1(T17N) was a gift from Klaus Hahn (Addgene plasmid # 22784; http://n2t.net/addgene:22784; RRID:Addgene_22784)^62^.

For construction of pK281, CyRFP excised with NheI and NotI from pCAG-CyRFP was blunted with SAP (Takara: 2660A) and Klenow (Takara: 2140A) and ligated into blunted SalI/EcoRV site of pK068 vectors (CAG-FRT-STOP-FRT-EGFP-ires-tTA-WPRE)^19^. For pK302, tagBFP excised with SalI/EcoRV from pK301 (CAG-loxP-STOP-loxP-tagBFP-ires-tTA-WPRE), for which tagBFP was cloned by PCR from pTagBFP-actin with primers KS130/KS131, was blunted with SAP and Klenow and ligated into blunted EcoRI site of pK038 vector (CAG-loxP-STOP-loxP-EGFP-ires-tTA-WPRE)^19^. For pK313, GCaMP6s was cloned by PCR from pK152 with primer pairs LW010/LW011 and was ligated into SalI/EcoRV site of pK068 vectors by using NEBuilder (New England BioLabs: E5520S). For pK326, we first constructed CAG-CreER^T2^. CreER^T2^ was cloned by PCR from pCreER^T2^ with primer pairs NN029/NN019 and ligated into SalI/NotI site of pK025 vector (CAG-turboRFP)^63^. Next, WPRE excised with NotI from pK068 was inserted into NotI site of CAG-CreER^T2^. Finally, ER^T2^ fragment cloned by PCR with primer pairs NN030/NN031 was inserted into SalI site of CAG-CreER^T2^-WPRE. For pK328, Rac1(T17N) was cloned by PCR from pCyPet-Rac1(T17N) with primer pairs SN088/SN089 and was ligated into site of pK029 vectors (CAG-loxP-STOP-loxP-RFP-ires-tTA-WPRE)^20^ by using NEBuilder.

Primer sequences were follows;

KS130: CTGTCGACATGAGCGAGCTGATTAAGGAGA; KS131: AGATATCTTAATTAAGCTTGTGCCCCAGTTTG; LW010: GAATAGGAACTTCATGAGATCTCGCCACCAT, LW011: TAACTCGATCTAGGATGCGGCCGCTCACTTC; NN019: CCCGCGGCCGCTCAAGCTGTGGCAGGGAAACCCT; NN029: GCTGTCGACAATTTACTGACCGTACACC; NN030: CCCGTCGACGCCACCATGGCTGGAGACATGAGAGC; NN031: ATTGTCGACAGCTGTGGCAGGGAAACCC; SN088: TATACGAAGTTATATGATGCAGGCCATCAAGT; SN089: ATCCTCGAGTCGCCGCTTACAACAGCAGGCAT.

### Drug administration

Clozapine-N-oxide (CNO) (Sigma: C0832, Tocris: 4936) prepared in saline was intraperitoneally injected (12 μg/g body weight)^64^. Lidocaine (AstraZeneca: Xylocaine) was subcutaneously injected (1%, <10 μL)^16^. Tamoxifen (Sigma: T5648) prepared in cone oil was intraperitoneally injected (50 μg/g)^65^.

### Craniotomy for *in vivo* imaging

Cranial window for *in vivo* imaging was made as described^38^. Mice were anesthetized with isoflurane (DS Pharma Animal Health, Zoetis). The skin over the right hemisphere was removed using scissors to expose the skull, and Vetbond (3M: 1469) was applied to seal the incision. Whisker-related cortical area was detected with increases of GCaMP fluorescence induced by whisker stimulation. A small piece of bone covering labeled neurons was removed with a sterilized razor blade (Feather: FA-10) leaving the dura intact. Gelfoam (Pfizer) was used to stop bleeding as necessary. Cortex buffer (125 mM NaCl, 5 mM KCl, 10 mM glucose, 10 mM Hepes, 2 mM CaCl_2_, and 2 mM MgSO_4_; pH 7.4)^66^ was applied during opening the skull. The custom-made titanium bar (T and I)^38^ was glued to the skull near the window. A 2.5 mm-diameter round cover glass (Matsunami: custom-made) was applied onto the exposed brain with 1% low melting point agarose (Sigma: A9793-100) in cortex buffer. The dental cement (GC Corporation: Unifast III) was applied to seal the cover glass and the titanium bar. For analgesic and anti-inflammation, carprofen (5 mg/kg, prepared in saline, Zoetis: Rimadyl) was subcutaneously injected.

### *In vivo* two-photon calcium imaging

*In vivo* two-photon calcium imaging was performed under an unanesthetized condition. A heater was used to keep pups warm. Time-lapse images (512 × 512 pixels, 12 bits) were obtained at 1 Hz using a two-photon microscope (Zeiss: LSM 7MP) with W Plan-Apochromat 20×/1.0 DIC objective lens (Zeiss). Mai Tai eHP DeepSee titanium-sapphire laser (Spectra-Physics) running at 920 nm or 940 nm was used. Fluorescent proteins were simultaneously excited, and emitted fluorescence was filtered (500−550 nm for green and 575−620 nm for red) and detected with LSM BiG detectors (Zeiss). Higher signal-to-noise ratio images were obtained with 940 nm or 960 nm wavelength laser and a slower scan speed under isoflurane anesthesia to visualize the barrel position in TCA-RFP Tg mice. Histological analyses after the end of *in vivo* imaging confirmed that imaged area was located within the (prospective) large barrel field of the primary somatosensory cortex.

### Histology

Mice were decapitated, and brains were fixed with 4% paraformaldehyde (PFA) in 0.1M PB at 4 °C for 1–2 days. Right hemispheres were flattened and transferred to 2% PFA/30% sucrose in 0.1M PB and kept at 4°C for 1–2 days. Tangential slices (100 μm thick) were obtained with a ROM-380 freezing microtome (YAMATO: REM-710) and mounted with Anti-fade Mounting Medium^67^. Slices which were not derived from TCA-GFP or TCA-RFP Tg were stained with anti-VGluT2 antibody (SYSY: 135403, 1:1000) and Alexa 647 goat anti-rabbit IgG (Invitrogen: A21244, 1:1000) and/or DAPI (Roche: 10236276001, 2 μg/mL) before mounting to visualize barrel map. Images were acquired by a confocal microscope (Leica: TCS SP5). For quantitative analyses of dendritic spines, spiny stellate neurons located at the edge of large barrels (α–δ, A–E arc 1–4) were chosen and their basal dendritic segments located inside the barrel were used. Spiny stellate neurons and barrel edges were identified as previously described^20^. Confocal images were taken with 40×/0.85 objective lens, 1.2 μm optical sectioning for measuring dendritic length, and with 63×/1.3 objective lens, 2× zoom, 0.1 μm optical sectioning for spine count. Dendrites were cut into segments by a series of concentric circles from 10 to 70 μm radius with 10 μm interval. Segments which contained dendritic tip and those which were highly overlapped with other dendrites were excluded from spine count analysis. Filaments were classified as follows. If length > 5 μm: Dendritic branch. Else if head width > 0.6 μm: Mushroom. Else if length > 2 μm: Filopodium. Else: Thin^68,69^. Dendritic lengths and spine counts were measured on 2D images with using 3D images as references.

### Slice electrophysiology

#### Whole-cell recording

For naïve P9 and P11, non-labeled mice were used. For DN-Rac1 and Tam control, L4 glutamatergic neurons were sparsely labeled by the IUE-based Supernova-RFP transfection. Oblique coronal slices of barrel cortex (300 µm thick) were prepared from P9 or P11 mice under deep anesthesia with isoflurane and kept in a normal artificial cerebrospinal fluid containing the following: 126 mM NaCl, 3 mM KCl, 1.3 mM MgSO_4_, 2.4 mM CaCl_2_, 1.2 mM NaH_2_PO_4_, 26 mM NaHCO_3_, and 10 mM glucose, saturated with 95% O_2_ and 5% CO_2_. Non-labeled neurons (for naïve P9 and P11) or RFP-positive neurons (for DN-Rac1 and Tam control at P11) in the barrel-cortex L4 were targeted by patch pipettes for whole-cell recordings under fluorescent and infrared differential interference contrast optics (Olympus: BX51). The patch pipettes were filled with an internal solution containing the following: 130 mM K-gluconate, 8 mM KCl, 1 mM MgCl_2_, 0.6 mM EGTA, 10 mM HEPES, 3 mM MgATP, 0.5 mM Na_2_GTP, 10 mM Na-phosphocreatine, and 0.2% biocytin (pH 7.3 with KOH). To record mEPSCs, tetrodotoxin (1 μM, Abcam) and SR95531 (100 μM, Abcam) were added to block action potentials and GABA_A_ receptors, respectively. The membrane potentials of recorded neurons were held at −75 mV. Whole-cell recordings were performed and sampled using Axon Multiclamp 700B amplifier, Digidata 1440A and pCLAMP10 software (Molecular Devices). Signals were sampled at 20 kHz. We selected cells with a high seal resistance (> 1 GΩ) and a low series resistance < 30 MΩ. mEPSCs were detected and analyzed using MiniAnalysis software (Synaptosoft).

#### Post-hoc morphological analysis

After electrophysiological recording, the slices were fixed with PFA in 0.1M PB overnight at 4°C for biocytin staining. To visualize recorded neurons, slices were incubated with streptavidin conjugated to Alexa 488 (1:1000; Life Technologies) in 25 mM PBS containing 0.1% Triton X-100 overnight at 4°C. Spiny stellate neurons were distinguished from other neurons by the spherical soma and the absence of a prominent apical dendrite^70^.

### *In vivo* image processing and quantification

#### Pre-processing

Green channel was extracted and Gaussian filter (sigma = 10 px for the active contour analysis, 2 px for the others) was applied on Fiji/ImageJ ver. 1.52p^71^.

#### ROI setting

∼15 μm × 15 μm rectangular areas were selected. In sparse labeling experiments, ROIs were basically put on cell bodies, but some ROIs were put nearby the center of the cell bodies to avoid signal saturation which can be the cause of failure of signal detection.

#### Signal detection and processing

Intensity of the ROI was the mean of the pixels in the ROI. Raw calcium signals for each ROI, F(t), were converted to represent changes from baseline level, ΔF/F(t) defined as (F(t) – F_0_)/F_0_, where F_0_ was the time-averaged intensity at each ROI. ΔF/F(t) > 1 was counted as neuronal firing. The time period from when ΔF/F(t) became over the threshold to when ΔF/F(t) became lower than the threshold was defined as one firing events.

#### Movie

For **Movs.1–3**, F_0_ was subtracted from Gaussian filtered images, and brightness was adjusted for visualization. Movie is 10 times faster than the real time.

#### Active contour drawing

Images were binarized based on ΔF/F(t) threshold, and the area of the active contour was measured using Fiji/ImageJ. If a contour protruded from the view field, it was excluded from the analysis. When a contour was observed in the same position over multiple timeframes, only the contour on the initial timeframe was used for the analysis.

#### Synchronized event detection

Monte Carlo simulation was applied to infer the chance rate of ROI synchronization. For each ROI, firing periods and non-firing periods were calculated and they were randomly shuffled. After all ROIs underwent random shuffling process, the number of ROIs fired together was counted. This process was repeated 1,000 times, and the distribution of active% in random firing was estimated as the proportion of the replications. 99th percentile of this distribution was used as the chance rate (p < 0.01). The time period in which the number of fired ROIs were over the chance rate was considered as a synchronized event.

### Statistics and computing

Fiji/ImageJ ver. 1.52p^71^ and custom-written scripts in Python 3.6.8 (Python Software Foundation) with its additional packages Numpy 1.16.4^72,73^, Scipy 1.2.1^74^, Matplotlib 3.1.0^75^, Pandas 0.24.2^76^, OpenCV 3.3.1^77^ were used to data analysis and visualization. Two-tailed Welch’s t-test was used to test the differences among means unless otherwise noted. The asterisks and pound symbols in the figures indicate the following: */# p < 0.05, **/## p < 0.01, and ***/### p < 0.001. p < 0.05 was considered statistically significant. g indicates Hedge’s g. In box plots, upper and lower limits of box represent 75th and 25th percentile, crosses represent mean, horizontal lines represent median, upper and lower whiskers represent maximum and minimum within 1.5 interquartile range, and observations beyond the whisker range were marked with open circles as outliers. Sample size was described in Results. Representative examples of each figure were chosen from the following number of mice; **Fig. 1**: 5, 3, and 2 mice at P1, P3, and P5, respectively; **Fig. 2**: 2 and 6 mice at P5 and P9, respectively; **Fig. 3**: 3 and 4 mice at P9 and P11−12, respectively; **Fig. 6**: 4 and 5 mice for Tam control and DN-Rac1, respectively. The following timeframes were used for each analysis; **Figs. 1d, 1e, 2, 3**: 240 frames (for **Fig. 3** P11, 3mice: 180 frames); **Figs. 1h, 4, 6**: 300 frames.

## Data and code availability

All data that support the conclusions and computational code used in the study will be available upon manuscript publication.

## Acknowledgements

We thank Pierre Chambon for the pCre-ER^T2^ vector; Takuya Sato, Minako Kanbayashi, and Satoko Kouyama for technical assistance; Naoki Nakagawa and Luwei Wang for helping plasmid construction; and Iwasato laboratory members for critical reading of the manuscript and stimulating discussions. This work was supported by KAKENHI (16H06460) to Y.Y. and KAKENHI (16H06459 and 20H03346) to T.I.

## Author contributions

S.N. and T.I. conceptualized the experimental design. Y. Y. and M.T. performed electrophysiological experiments and analyses. H.M. contributed to setting up the *in vivo* calcium imaging. S.N. performed most experiments, analyses and data visualization. S.N. and T.I. wrote the manuscript. All authors reviewed the manuscript.

---

**Movie 1**. Spontaneous activity in the barrel cortex at P1. L4 neurons were densely labeled with CAG-GCaMP6s.

**Movie 2**. Spontaneous activity in the barrel cortex at P9. L4 neurons were densely labeled with CAG-GCaMP6s.

**Movie 3**. Spontaneous activity in the barrel cortex of Tam control and DN-Rac1 mice at P11. L4 neurons were sparsely labeled with Supernova-GCaMP6s.

